# Molecularly distinct astrocyte subpopulations spatially pattern the adult mouse brain

**DOI:** 10.1101/317503

**Authors:** Mykhailo Y. Batiuk, Araks Martirosyan, Thierry Voet, Chris P. Ponting, T. Grant Belgard, Matthew G. Holt

**Author notes:** To whom correspondence should be addressed: Dr. Matthew Holt, Herestraat 49, B-3000 Leuven, Belgium. These authors contributed equally to this work (Mykhailo Y. Batiuk, Araks Martirosyan). Present address: MRC Human Genetics Unit, University of Edinburgh, UK. Present address: Verge Genomics, 42A Dore Street, San Francisco, CA 94103, USA.

## Abstract

Astrocytes are a numerous cell type of the central nervous system. They perform many important functions in synapse formation and maintenance, control of local homeostasis and modulation of synaptic transmission. However, the degree to which specialist astrocyte subtypes fulfil these specific tasks is currently unclear. Here we use single cell transcriptomics to demonstrate that astrocytes, even those lying within the same region of adult mouse brain, show distinct molecular and spatial profiles, with profound implications for local CNS functions, including axon guidance, synaptogenesis and synaptic transmission.

---

Astrocytes have traditionally been considered to be a homogeneous set of cells. However, recent reports suggest that astrocytes are, instead, heterogeneous, with considerable variation in morphology, physiology and function across brain regions (comprehensively reviewed in^1–3^). Nevertheless, it is unknown whether specialized astrocyte subsets perform distinct functions, between and within CNS regions. If so, this could have profound consequences for local neuronal function in both the healthy and diseased brain^1, 2^.

We sought to obtain an in-depth and unbiased perspective on astrocyte heterogeneity, both within and between major brain regions, using single cell mRNA sequencing. To accurately distinguish between transcriptional programs associated with development and those underlying functional specialization in the mature brain^4^, we targeted astrocytes isolated from adult (post-natal day 56) mouse brain. For cell isolation from both hippocampus and cortex (selected for their well-studied anatomy, physiology, and broad disease relevance), we used a FACS-based protocol, based on staining with the ubiquitous astrocyte marker ATP1B2^5^ (Online Methods; Supplementary Text; Supplementary Figures 1-5).

In total, we isolated 2,976 cells and prepared individual cDNA libraries using a modified SMART-Seq2 approach (Online Methods; Supplementary Text; Supplementary Figure 6), chosen for its sensitivity and full-length transcript coverage^6^. Libraries were sequenced using Illumina sequencing and results were quality controlled using standard criteria (Supplementary Figures 7-8). Following removal of low quality data, 2,031 libraries remained, with an average of 2,148 genes detected per cell.

Seurat^7^ based clustering on these libraries revealed 6 distinct clusters of higher order cell types, based on expression of known marker genes (Figure 1a,b; Supplementary Figure 9; Supplementary Table 1). Following removal of contaminating cell types, 1,811 astrocytes were reclustered using Seurat, based on 886 highly variable genes. This led to the identification of 5 distinct <u>**A**</u>strocyte <u>**S**</u>ub<u>**T**</u>ypes (AST1-5) each distinguished by gene expression fingerprints (Figure 1c,d; Supplementary Figures 10-14), which reveal the most subtle characterization of these brain regions reported to date^8^.

**Figure 1.**
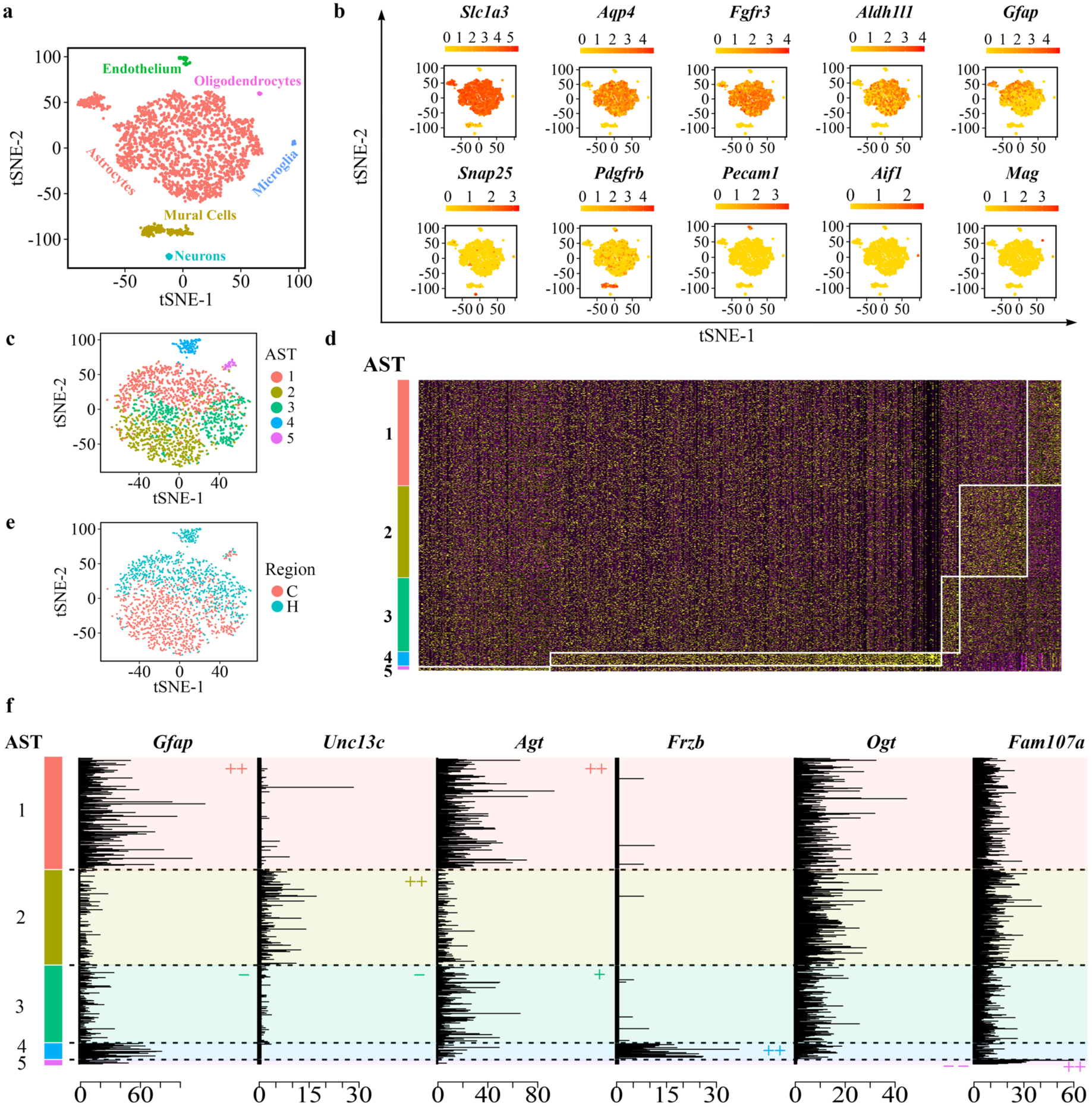
Computational identification of multiple astrocyte subtypes in adult mouse cortex and hippocampus. (a) Visualization of the major higher order cell types identified by Seurat, using t-distributed stochastic neighbor embedding (tSNE) plots. Each dot represents a single cell. Different cell types were assigned according to expression of cell type specific marker genes and were labeled with different colors. (b) tSNE plots of higher order cell types highlighted for expression of well-known astrocyte markers (*Slc1a3*, *Aqp4*, *Fgfr3*, *Aldh1l1* and *Gfap*). Markers of other cell types (*Snap25*, neurons; *Pdgfrb*, mural cells; *Pecam1*, endothelial cells; *Aif1*, microglia; *Mag*, oligodendrocytes) cluster separately from astrocytes. Normalized gene expression data is shown in ln-scale. (c) Seurat-based clustering of astrocyte data identified 5 distinct astrocytes subtypes (AST), marked with a distinct color code. Data are presented in tSNE plots. (d) Hierarchically clustered gene expression heatmap of subtype overexpressed genes across 1,811 astrocytes. Rows correspond to cells, columns to genes. Yellow, high gene expression level; black, no gene expression. Normalized gene expression data is shown in ln-scale. (e) Distribution of astrocytes derived from hippocampus (H) and cortex (C). (f) Astrocyte subtype-specific marker genes selected for *in situ* hybridization experiments, based on normalized expression data. Positive markers for each subtype are indicated with “+” or “++” depending on expression level. In contrast, negative markers are labelled “-” or “--”.

Astrocyte subtypes constitute distinct proportions of the total cell population (Supplementary Table 2), with clear separation of two astrocyte subtypes between cortex and hippocampus (Figure 1e; Supplementary Table 2) providing the first indications of functional specialization between regions. Extensive controls excluded clustering driven by batch effects arising during the preparation of sequencing libraries (Supplementary Figures 15-18).

Unique sets of marker genes imply distinctive subtype specializations. Gene enrichment and functional annotation^9^ (Supplementary Table 3), as well as manual curation of genes^10^ (Supplementary Tables 4-6), identified transcription factors specific to each subtype, which implies sustained maintenance of diverse underlying gene expression programs. Genes relating to neurite outgrowth and synaptogenesis exhibited contrasting expression profiles across these astrocyte subtypes, even for those residing within the same brain region (such as ASTs 2 and 3, Supplementary Table 2). These results are indicative of synaptic micro-patterning by astrocyte subtypes, as exemplified by local spinal cord wiring directed by a population of *Sema3a* positive astrocytes^11^, and suggest that this represents a general CNS phenomenon rather than an exception. In particular, AST2 appears to be specialized for local production of active thyroid hormone (T3) which exerts powerful effects on CNS function and synapse development, including regulating production of the key synaptogenic molecule cholesterol^12,13^. Furthermore, AST2 overexpresses genes linked to glutamatergic neurotransmission, even including those involved in the mobilization of the potent NMDA receptor co-agonist D-serine^14^, indicating that specific wiring connectivity reflects distinct synaptic functions.

Functional specializations of astrocyte subtypes could reflect their location within tissue niches and their influence on the local CNS environment. Each astrocyte subtype was identified with a signature of multiple genes (Figure 1f; Supplementary Table 7).

Using high resolution mapping of these markers by multiplexed *in situ* hybridization (ISH), the five astrocyte subtypes were revealed to occupy distinct cortical layers and hippocampal locations (Figures 2-4; Supplementary Figure 19), consistent with low resolution data on these markers from single probe staining deposited in the GENSAT database^15^ and Allen Mouse Brain Atlas^16^ (Supplementary Figures 20 and 21). AST1 was found preferentially in the adult derivatives of the marginal zone facing the pia and, surprisingly, around the hippocampal fissure. AST2 was mostly present in cortical layers 2-5. In contrast, AST3 was widely distributed throughout cortical layers 1-6, as well as across the majority of the hippocampus. AST4 was located in the subgranular zone of the dentate gyrus and showed co-staining with known stem cell markers consistent with it representing a population of neural stem cells. Finally, AST5 was found as rare, isolated cells mostly within layers 1 and 6 of the cortex and the stratum lacunosum-moleculare of the hippocampus.

**Figure 2.**
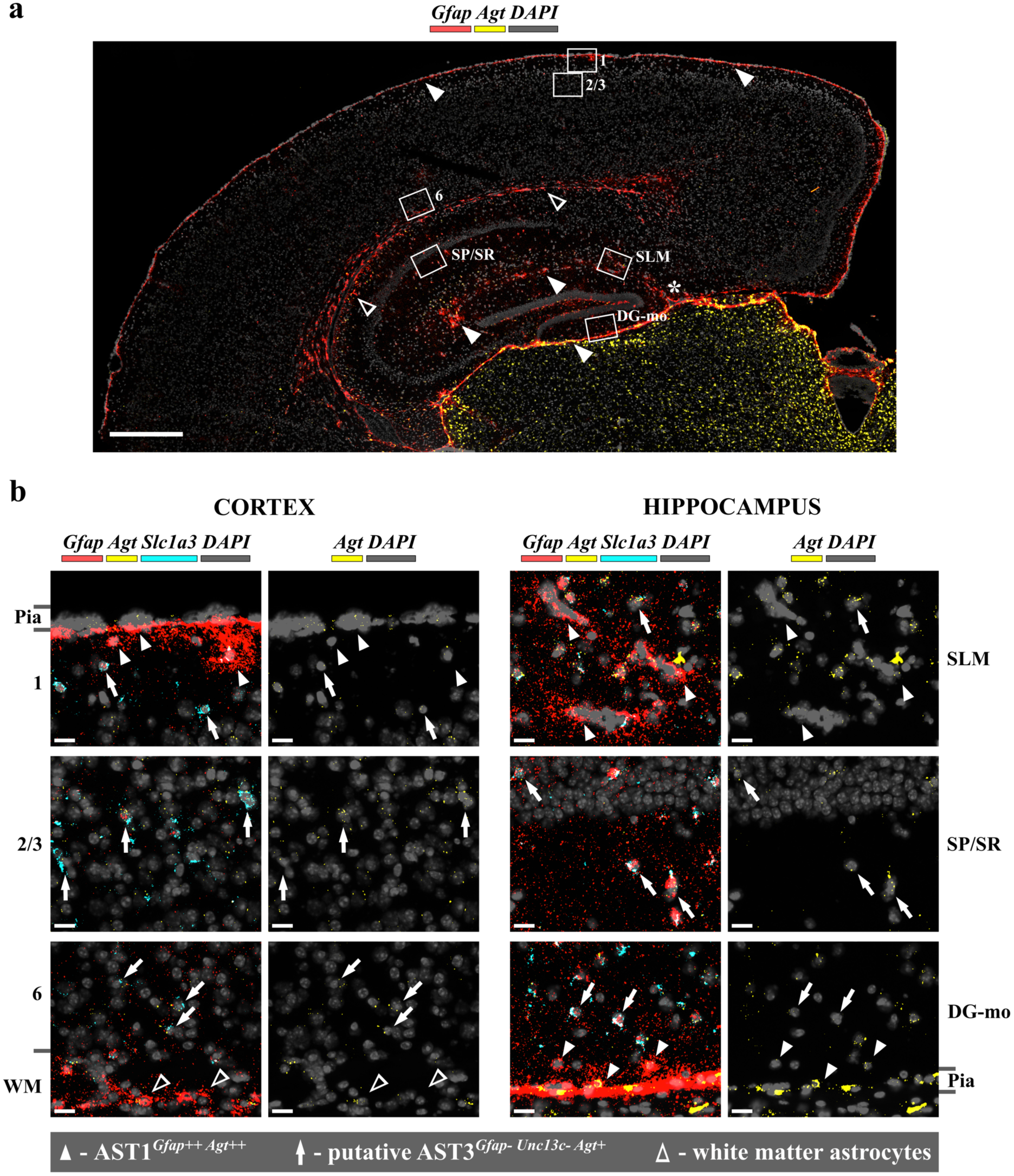
Differential patterning of AST1 and AST3 in adult mouse brain mapped with *in situ* hybridization. (a) Overview of a coronal section of adult mouse brain stained for AST1^*Gfap++ Agt++*^. AST1 was detected in the subpial area in cortex (layer 1) and stratum lacunosum-moleculare (SLM) of the hippocampus (closed arrowheads). White matter astrocytes also highly express *Gfap* (open arrowheads), although this region was removed during tissue preparation. Asterisk (*) marks the hippocampal fissure. Boxes mark regions represented in high magnification in (b). Scale bar, 500 μm. (b) High magnification images highlighting the position of AST1 astrocytes (defined by co-expression of *Slc1a3*) in cortex (left) and hippocampus (right) (closed arrowheads). Different cortical layers are indicated numerically. Hippocampal regions are indicated as SLM, SP/SR (stratum pyramidale/stratum radiatum) and DG-mo (molecular layer of dentate gyrus). Position of the white matter is indicated (WM). AST1 was mapped primarily to the subpial area in layer 1 of cortex, the subpial area in hippocampus and the SLM around the hippocampal fissure. Astrocytes showing expression of *Agt* and low expression of *Gfap* in cortical layers 1-6, as well as the majority of zones in the hippocampus, are likely to represent AST3^*Gfap-Unc13c-Agt+*^ (ar rows). Scale bars, 20 μm.

**Figure 3.**
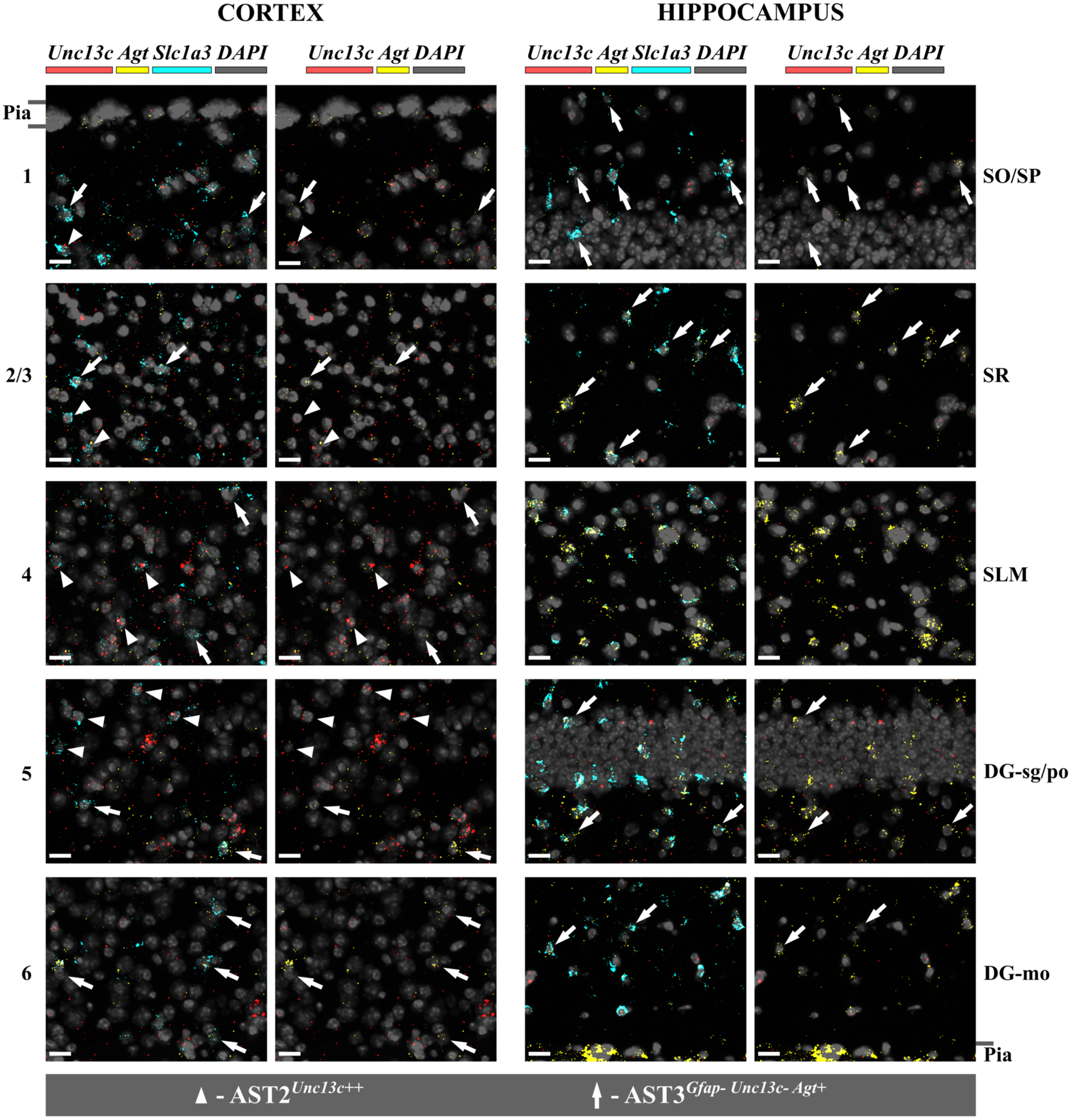
Differential patterning of AST2 and AST3 in adult mouse brain mapped with *in situ* hybridization. High magnification images highlighting the position of astrocytes (defined by co-expression of *Slc1a3*) in cortex (left) and hippocampus (right). Different cortical layers are indicated numerically. Hippocampal regions are indicated as SO (stratum oriens), SP (stratum pyramidale), stratum radiatum (SR) and stratum lacunosum-moleculare (SLM). Specific regions of the dentate gyrus including the stratum granulosum/polymorphic (DG-sg/po) and molecular layer (DG-mo) are marked. AST2^*Unc13c++*^ was detected primarily in cortical layers 2-5 (arrowheads). AST3^*Gfap-Unc13c-Agt+*^ was detected throughout layers 1-6 of cortex and SO, SP, SR, DG-sg/po and Dg-mo (arrows). Scale bars, 20 μm.

**Figure 4.**
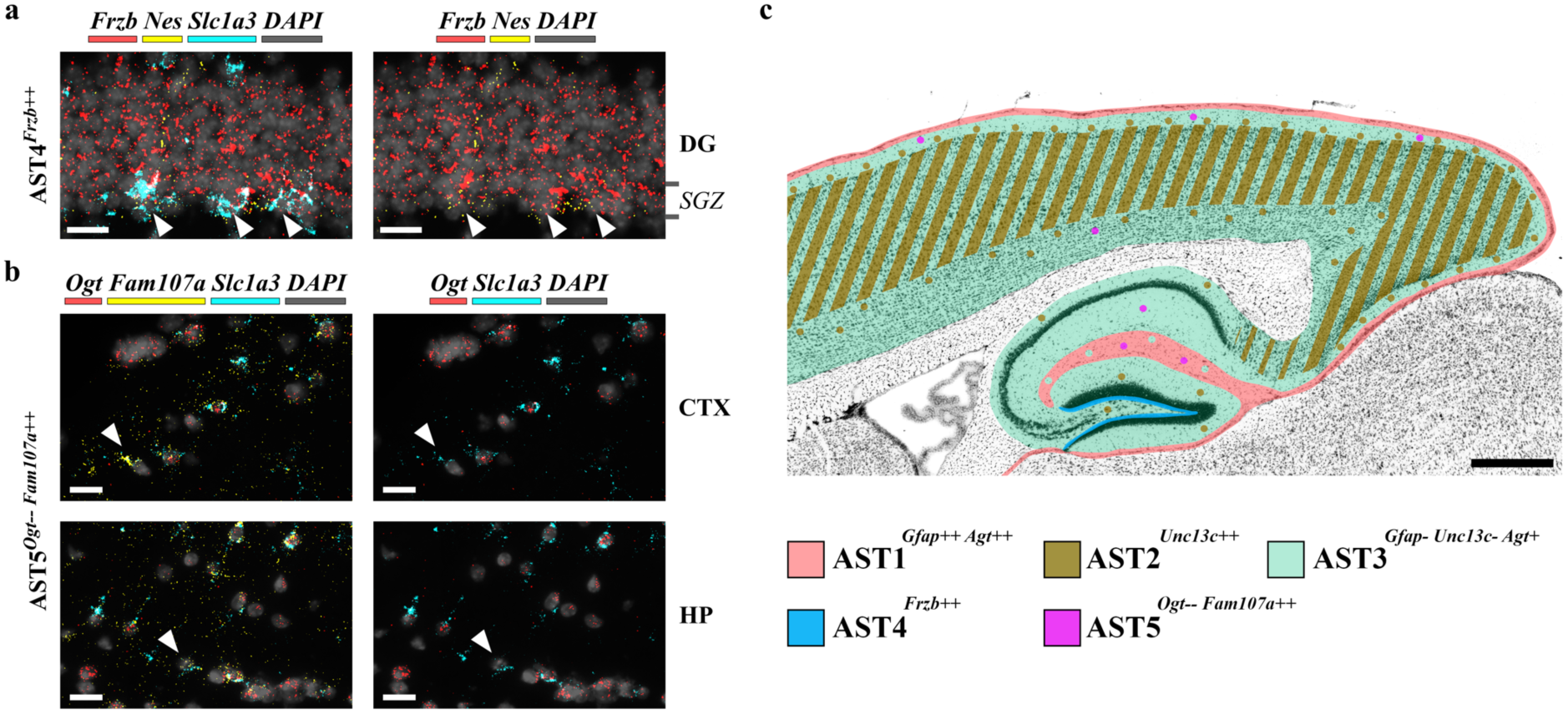
Differential patterning of AST4 and AST5 in adult mouse brain mapped with *in situ* hybridization and a schematic overview of AST positioning. (a) High magnification images highlighting the position of AST4^*Frzb++*^ astrocytes in brain (defined by co-expression of *Slc1a3*). AST4 was detected in the subgranular zone (SGZ) of the dentate gyrus (DG) in hippocampus, a known site for adult neurogenesis. Co-staining with the known stem cell marker *Nestin (Nes)* putatively identifies this population as containing adult stem cells. Scale bars, 20 μm. (b) High magnification images highlighting the position of AST5^*Ogt-- Fam107a++*^ astrocytes (defined by co-expression of *Slc1a3*). Cells were sparsely located through the cortex (CTX) and hippocampus (HP) and are marked with arrowheads to aid identification. Representative images from layer 1 of cortex and the stratum lacunosum-moleculare of the hippocampus are shown. Scale bars, 20 μm. (c) Summary of astrocyte subtype positions in adult mouse cortex and hippocampus. Positions are marked on a representative sagittal section of adult mouse brain (adapted from the Allen Mouse Brain Atlas). Subtypes are color-coded. Scale bar, 500 μm.

The observation of unique spatial patterning of astrocyte subtypes suggests several intriguing possibilities, many of which have been widely debated in the literature, but which have not had the benefit of the experimental resolution afforded by single cell methodologies. For example, the widespread distribution of ASTs 1, 3 and 5 across cortex and hippocampus hints at a shared developmental origin from embryonic pallium^17^.(Supplementary Figure 22). By contrast, in both mouse brain and spinal cord, evidence exists for astrocyte generation from distinct progenitors, with developmental patterning irreversibly fixed to the site of progenitor origin^18^. Superficially, this appears incompatible with the cortical co-location of ASTs 2 and 3. However, regional intermixing of astrocyte subtypes, each defined by a complex molecular signature of multiple genes, may well be explained by origination from a common progenitor, followed by functional refinement dependent on local neuronal activity^19^. Finally, the concepts of migration and local maturation^20^ may also explain the transcriptomic similarities between the rarest astrocyte subtype, AST5, and AST4 - suggesting that AST5 represents an intermediate along a developmental pathway starting with stem cells and ending with differentiated astrocytes.

In summary, we have provided detailed evidence for an unprecedented degree of inter- and intraregional heterogeneity of astrocytes and have shown, for the first time, both distinct cortical layering and hippocampal compartmentalization of molecularly distinct astrocyte subtypes. Although our appreciation of such heterogeneity is likely only to rise, as single cell transcriptomes of additional brain regions are sequenced, our work provides a highly resolved roadmap for future investigations of astrocyte development and function. It generates testable hypotheses relating to the properties of astrocyte subtypes that, when allied with the increasing power of intersectional genetics^21^, will ultimately allow their effects on neuronal form and operation to be elucidated. Such information should prove invaluable to our overall understanding of CNS development and function.

## Methods

Detailed methods are available in the online version of the paper.

## Data availability

The full list of markers and the sequencing count table (not normalized) are provided online as Supplementary Files. Raw data will be made publicly available through the GEO database (number to be provided).

## Author contributions

MGH conceived and directed the project. MYB isolated cells, performed FACS experiments, developed the modified Smart-Seq2 protocol, prepared single cell cDNA libraries and performed the ISH experiments. MYB also performed gene function analysis using David and Uniprot. AM performed Seurat-based analysis under the guidance of TGB and CPP. TV provided access to equipment. MGH wrote the final manuscript, with input from all co-workers, particularly MYB and CPP. All authors approved submission.

## Acknowledgements

MGH grateful acknowledges support from the European Research Council (Starting Grant 281961), Fonds Wetenschappelijk Onderzoek (FWO) (G066715N) and VIB Institutional Support. TGB and CPP were supported by the Medical Research Council. TGB was additionally supported by the Human Brain Project (European Union Seventh Framework Programme (FP7/2007-2013) Grant Agreement 604102). TV is supported by Fonds Wetenschappelijk Onderzoek (FWO) (G066715N).

The authors wish to express thanks to the KU Leuven FACS core, LIMONE imaging unit, VIB Nucleomics core, Mark Fiers (VIB Bioinformatics Unit) and Ligia Mateiu (Voet lab) for help at various stages during this project.

